# Intensity matching in cuttlefish

**DOI:** 10.1101/018176

**Authors:** Andres Laan

## Abstract

For efficient background matching it is essential that animals closely match a set of salient visual statistics of their visual surroundings. The mean intensity of a background is a key statistic, because it can be estimated across a large range of viewing distances by a simple computation. We investigated how the dynamic neuromuscular camouflage system of the cuttlefish *Sepia offcinalis* responds to changes in the mean background intensity of uniform backgrounds. We find that cuttlefish adapt their body intensity in response to variations in the mean background intensity, yet show biases in their body intensity beyond what can be predicted from the limited dynamic range of their camouflage system. On sandy backgrounds of various reflectance values their uniform body patterns maintain a constant yellow hue. This color constancy may represent an example of a color prior in a colorblind animal because a yellow body color would be the optimal hue for camouflage on sands typically encountered in their natural environment. Cuttlefish adapt their appearance to the background via a dynamic process composed of a complex mixture of intensity transients spanning timescales from the subsecond to the minute range. In very young animals camouflaging on dark sands the masquerade strategy is preferred over background matching. Masquerade is implemented by combining partial background matching with frequent expression of disruptive components. We furthermore provide an objective definition of disruptive components using hierarchical clustering and automated image analysis thus highlighting the role of chromatophore activity correlations in structuring the motor output of *S.officinalis*.

## Introduction

The soft-bodied cuttlefish uses camouflage to avoid detection by its predators (1). Two features present considerable challenges to a camouflaging cuttlefish- the varied and complex nature of its visual surroundings and the keen eyesight and diversity of visual systems among its many predators (2). Many animals facing less sophisticated visual predators or less diverse visual surroundings have evolved simple yet effective heuristic solutions for camouflage. Examples include the stripes of zebras, which are effective at deceiving the visual system of tse-tse flies (3) or the white fur of artic hares (4), which enables a visual blending into its monotone artic surroundings.

That the camouflage system of the cuttlefish is much more sophisticated can be readily appreciated by an examination of its physical structure (1). The body of a cuttlefish is tiled with millions of tiny pigment sacks called chromatophores. Each chromatophore is surrounded by a set of radial muscles whose expansion level is under direct neuromuscular control. The activity of the radial muscles determines the size of the chromatophore and allows regulation of the local intensity of a patch of skin. The dense tiling of chromatophores on the skin provides the cuttlefish with physical machinery that rivals modern printing and telecommunications technology in its capabilities.

Yet such sophisticated camouflage machinery would be ineffective without a concomitantly sophisticated control system regulating the generation of camouflage patterns. Such a control system must implement a sensorymotor transformation from the visual surroundings to an appropriate body pattern. A collective body of experimental work has used the camouflage response as a sensorymotor assay to determine the many parameters of its visual environment to which cuttlefish are sensitive. These parameters include contrast (5), the presence of edges (6), intensity (7), polarization (8), image frequency content (6) and more complex configural features such as sensitivity to illusory contours (9), but surprisingly exclude information regarding color (10). In parallel, behavioral work has identified large-scale units of chromatophores commonly called components that form recognizable units from which body patterns are assembled (11). What is required to further refine our understanding of cephalopod camouflage is a comprehensive set of quantitative rules, which define the transformation of visual features into cuttlefish body patterns.

When considering background matching on visual textures, many authors have expressed hope that the theory of image statistics will provide a principled way forward towards a normative theory of camouflage (12). In the early 1970s Bela Julez investigated various statistics of images beginning with the mean, variance and autocorrelation and put forward a conjecture that two textures which have matching values for a set of statistics will be perceptually indistinguishable (13). While the Julez conjecture remains unproven and the necessary set of statistics unknown, results from computer vision (14) indicate that when images are analyzed with sets of local oriented filters (commonly called wavelets, similar in appearance to the receptive fields of cortical visual neurons found in mammalian visual cortex (15) but also across the animal kingdom (16)) and the resulting analysis coefficients are summarized in a set of a few hundred statistics, the resulting numbers can be effectively used to synthesize arbitrary textures to generate a very good perceptual match between template and synthetic images.

The systematic method of pattern generation by a cuttlefish might therefore involve the calculation of the salient statistics of its visual surroundings followed by choosing from among its many body patterns the one that minimizes the discrepancy between surround and body pattern statistics or defaulting to a body pattern such as disruptive (which uses the eponymous mode of camouflage rather than background matching) in case no good match is found.

Even with this rather specific hypothesis about the method of cuttlefish camouflage, we are left with many ambiguities the most important of which is the question regarding the necessary set of statistics. It is not even known whether the set of statistics is unique for any given wavelet transform and little is known about the effects that a change of basis has on the necessary set of statistics.

In an effort to find a suitable starting point for investigating a normative theory of camouflage under all these uncertainties, we reasoned that the mean intensity of a texture would be the most relevant statistic for two reasons. First, cuttlefish are likely to be viewed by predators from a large variety of distances. The angular resolution of the visual system is limited due to blurring by the optics of the eye and the limit imposed by the finite size of photoreceptor angular spacing. Thus, increasing viewing distances would progressively degrade access to information contained within the high frequency components of an image. Information about the mean is contained within the lowest frequency bands and thus could be estimated from the greatest range of viewing distances. Secondly, the mean is a statistic that should be easy to estimate in most predator visual systems as its estimation involves a simple summation of photoreceptor activities within the appropriate retinal region.

We thus chose to investigate whether and how well cuttlefish match this most important statistic by placing them on a series of seven backgrounds of uniform sands with increasing intensities varying from black to white. We found that cuttlefish varied their mean intensity as function of background intensity, but showed a series of systematic biases not dictated by the capabilities of their motor system. They were consistently brighter than the background at low background intensities and dimmer than background on the brightest backgrounds. Despite being exposed predominantly to grey backgrounds their color was a low saturation yellow at all intensities. Their adaptation to the background was a gradual process spanning several hundreds of seconds, despite their well-known capability to dramatically transform their appearance within less than a second. Finally, juvenile animals displayed an increasing activity of their disruptive components as background intensity decreased thus showing evidence of using a mode of camouflage different from their older conspecifics.

## Results

We prepared four sand backgrounds of uniform intensity (white, brown, grey and black), granularity and illumination to investigate animal responses to changes in mean background intensity. The intensity response of the animal to a background was characterized by calculating the mean intensity of the animal in 6 frames located 5,9,13,17,21 and 25 minutes from the start of the experiment. The animal response was then estimated as the average of those six measurements. The intensities were calculated in camera units of intensity (see methods) and we used the green intensity channel as the spectral sensitivity of that channel most closely matches the sensitivity of the cuttlefish visual pigment (the qualitative conclusions of the following analysis also held true when images were converted from RGB to greyscale prior to analysis).

For each background between 6 to 10 animals approximately three months in age were tested and the animal responses were averaged over the number of animals tested to calculate the population intensity for each background. The plot of population intensity versus background intensity clearly demonstrates that animals adapted their intensities to the intensities of their background (Figure 1A,B). Surprisingly, the animals displayed biases that were significantly different from background intensity on each of the four backgrounds tested. For the three darkest background animals showed patterns brighter than the background, whereas for the lightest backgrounds the animals were darker than the background.

**Figure 1:**
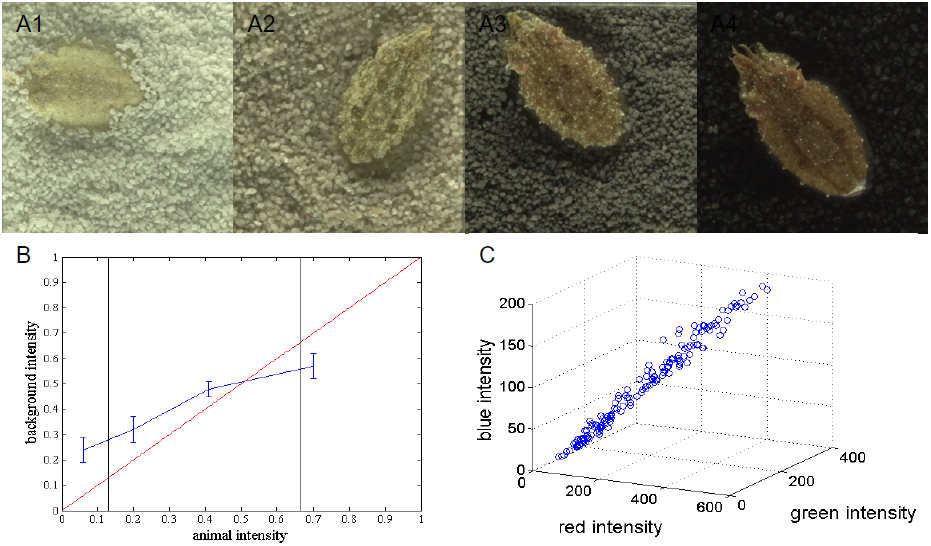
Biases in cuttlefish intensity matching. A1-4 Example animals camouflaging on white, brown, grey and black backgrounds respectively. B Summary graph of intensity matching. The blue curve shows the mean intensity of the population of animals for each of the four backgrounds, error bars show standard deviation. The red line illustrates unbiased intensity matching. The black bars delineate the intensity range within which unbiased background matching is possible given the dynamic range of the cuttlefish camouflage system. C Color structure of the animal. A plot of the mean intensities of 150 patterns in RGB space. The first principal component accounts for 99% of the variance showing that animals keep a constant yellow hue across the their full dynamic range of mean body intensities.

We first speculated that the limited dynamic range of the camouflage system might cause the biases on darkest (black) and brightest (white) backgrounds. We determined the brightest possible pattern of the camouflage system by exposing five animals to magnesium chloride, an anesthetic agent that causes a paling of the animals and is presumed to act as a muscle relaxant (17). The mean intensity of animals thus anesthetized was intermediate between the intensity of the white background and the population response on the white background. The difference between the anesthetized and white population response was statistically significant (p=0.007) demonstrating that the biases shown on the brightest backgrounds were only partly accounted for by the restricted dynamic range of the motor system. We estimated the dark limit of the chromatomotor system by finding the darkest individual pattern from the samples from which the population means were calculated. This response was similarly intermediate between the intensity of the black background and the population response on the black background demonstrating that motor system limitations again do not fully account for biases shown on the darkest backgrounds. For the intermediate intensity backgrounds from grey to light brown, the background intensity values lay within the dynamic range of the motor system, yet biases were still shown on these backgrounds.

Periods of motion have been reported to be correlated with reduced periods of camouflage (18). We hence automatically tracked all animals and excluded periods of motion from the analysis of the mean intensity (the first 5 minutes after introduction were excluded from analysis for reasons of comparability and stationarity, see below). Exclusion of periods of motion from intensity analysis gave mean animal intensity values which were only 3% lower than mean intensity calculated using the whole time period, but the difference between the two values was not statistically significant. We conclude that the biases found represent a robust experimental finding not explained either by animal motion or the restricted dynamic range of the motor system. The function and origin of these biases presently remains unknown.

In addition to biases in mean intensity, the animals also displayed systematic biases in their color. Cuttlefish possess two dominant color classes of chromatophores (yellow and black (1)). Because each color class can be controlled independently, the system has two degrees of freedom and the space of possible mean intensities in RGB space is expected to be two-dimensional. When we plotted the mean luminance of 154 different patterns in RGB space, we surprisingly found that most of the points fell onto a straight line (Figure 1C). Principal component analysis confirmed that a single dimension could account for 99% of the observed variance in the data. It thus appears that during intensity matching the activities of yellow and black chromatophores remain tightly correlated. The results of such a correlation is that the animals retain a low saturation yellow hue across the their full dynamic range of body intensities.

The dynamics of the chromatomotor system allow the animal to dramatically change its appearance within less than a second and such fast appearance transformations have indeed been observed in the context of interspecific threat signaling (1) and dynamic wave displays (19). Yet little is known about the timescales under which animals adapt to their environments in the context of camouflage. We measured the dynamics of intensity changes of our experimental animals for ten minutes following introduction to the experimental tank (3Hz sampling rate). The population as a whole adapted to the background with an exponential decay with a 100 second time constant (Figure 2A). At the level of individual animals, the time course of intensity changes was much more diverse and irregular (Figure 2C). Individual animals displayed intensity transients ranging from the subsecond to the minute timescale (Figure 2C,D).

**Figure 2:**
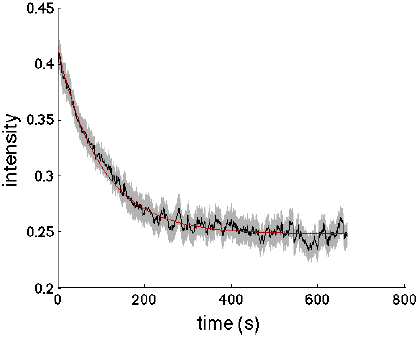
Timescales of intensity transients during adaptation to background. A Plot of the time course of the population intensity response on the two darkest backgrounds (black curve, grey region shows s.e.m). The mean intensity of an animal was sampled at 3Hz for 10 minutes following the introduction of the animal to the experimental tank lined with black or dark grey sand. Each time point represents the average over 16 animals. The population as a whole adapts to the background with an exponential decay depicted in red. Best fit value for the time constant is 100 seconds.

When we tested camouflage responses on four sands in younger animals around 1 month in age, we were able to confirm our findings of intensity biases (Figure 3A,B), which were qualitatively similar on all four backgrounds (a quantitative comparison of biases is not appropriate because the younger animals produced different classes of patterns). On black and dark grey backgrounds we surprisingly found that the animals frequently preferred to adopt a strongly disruptive rather than a uniform pattern. We found a background intensity dependent increase in the tendency of young animals to display disruptive patterns, which could also be quantified as an increase in energy in low frequency bands of the spectra (5) of young animals.

**Figure 3:**
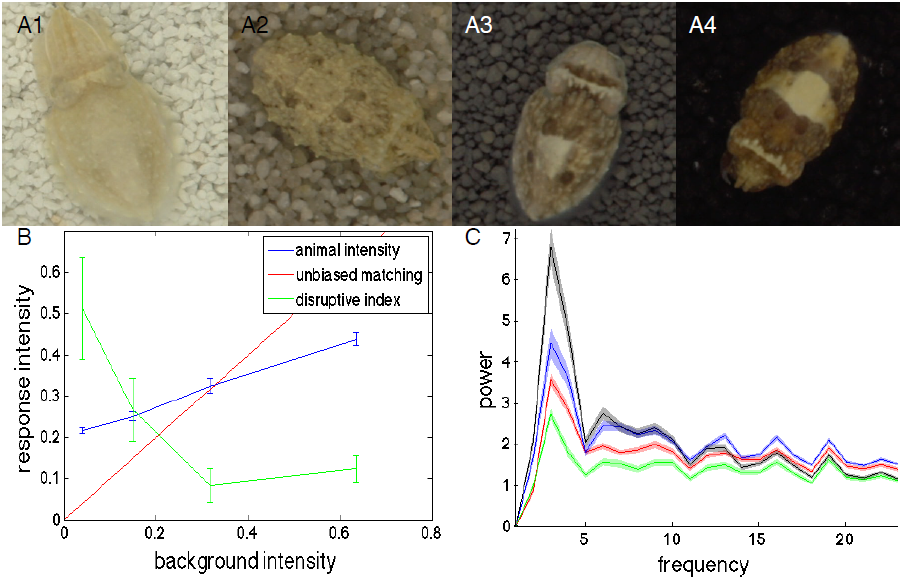
Differences in intensity matching between juveniles (1 month old) and young adults (2.5 month old) A1-4 Example juvenile animals camouflaging on white, light grey, dark grey and black sands. Note the disruptive patterning seen in the animals on dark grey and black sands. B Summary graph of intensity matching for juvenile animals. The red curve illustrates unbiased intensity matching, the blue curve illustrates the behavior exhibited by the animals. Note that juveniles like young adults adapt to the background intensity and show similar biases. The green curve plots the average disruptive index (calculated from visual annotation, varies between 0 and 1 for uniform and full disruptive respectively) as a function of background intensity. Darker backgrounds increase disruptive tendencies. C Quantification of body pattern differences on different sands is here demonstrated as an increase in disruptive band energies in the animals Fourier spectrum as the background intensity decreases (color-background correspondence: black-black, blue-dark grey, red-white, green-brown).

Despite being clearly identifiable visual elements on the skin of cuttlefish an objective definition of a disruptive component has been lacking in the literature. We utilized the tendency of young cuttlefish to express disruptive components to assemble a library of 170 disruptive patterns. We subsequently morphed all 170 images to conform to a common cuttlefish template and subjected the morphed image ensemble to hierarchical clustering (the head was excluded from analysis due to its frequent partial occlusion by the mantle edge). The resulting clusters clearly resembled the disruptive components and their edges (Figure 4). Because image segmentation can be based on both pixel intensity correlations and edges (20), we computed the edge density map over the image ensemble. The map of edge density showed clear density maxima around the borders of disruptive components. We thus propose that disruptive components can be defined as large territories of correlated chromatophore activity with sharp borders between the regions.

**Figure 4:**
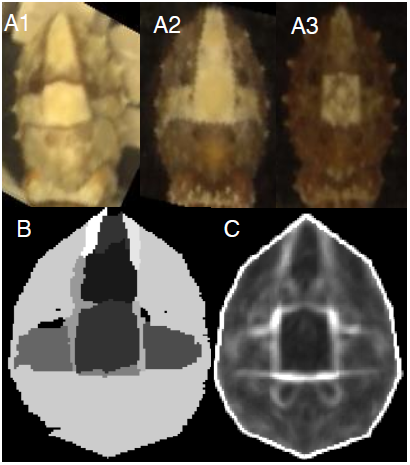
Automated extraction of disruptive patterns. A1-4 Four example animals showing patterns illustrating a uniform (A1) and three different kinds of disruptive (A2-4) body patterns. B Components extracted after 170 patterns were morphed to a common template and then analyzed by hierarchical clustering. The resultant components closely map on to disruptive components previously identified from behavioral observations. C The average edge density map on the body of a cuttlefish outlines the borders between the disruptive components. Two frontal eye spots are also visible.

## Discussion

We investigated how cuttlefish matched the mean intensity of uniform backgrounds. To our surprise, we found systematic biases in the intensity responses of cuttlefish with respect to both mean intensity and color. The biases in mean intensity were not caused by limited dynamic range or a motion-dependent break in camouflage. A finding of biases in intensity matching is not unique to cuttlefish (21). An adaptive pigmentation system must balance camouflage requirements against other physiological functions such as thermoregulation (22) or predation risk (21). It is presently unclear what if any functional purpose is served by intensity biases in cuttlefish.

In our experiments, we exposed cuttlefish to grey sands yet found that the animals retained a low saturation yellow hue across their entire observed body intensity dynamic range. On grey sands of varying intensity such a response represents a systematic color bias. In their natural habitat however, the situation is rather different. Cuttlefish face the challenge of having to control their body color for camouflage despite being colorblind (10). They must thus assume the color of their background based on evolutionary knowledge. In the natural environment of cuttlefish, yellow sands predominate as the typical substrate. The color biases they exhibit in our experiment might thus represent an evolutionarily adaptive prior knowledge about the predominant colors of sands in their natural habitats.

By developing automatic tracking system we were able to quantitatively monitor animal intensity as they adapted to dark backgrounds. Individual animals displayed considerable variation in their adaptation behavior and showed intensity transients spanning a 100-fold range of timescales. In contrast, the population as whole relaxed towards the steady state intensity in a regular exponential decay process. Exponential relaxations are characteristic of negative feedback control systems and stochastic Poisson processes. Some combinations of negative feedback based adaptation (23) and a stochastic stepwise relaxation process (24) may explain both the finding of irregularity in individual waveforms and regularity across the population. While cephalopods are able to adapt to backgrounds in the absence of visual feedback (1), the observation of exponential decay suggests that negative feedback may play a role in guiding camouflage responses under some conditions.

The camouflage strategies adopted by cuttlefish on uniform sands were age-dependent. While older animals employed uniform or weakly mottled patterns, young animals frequently produced disruptive patterns. The tendency of young animals to express disruptive components on uniform backgrounds seems at first sight counterproductive, because such components cause a sharp break of the visual statistics of an animal’s body from its uniform visual surroundings. Nor is our result explained by the fact that a typical sand grain size is larger relative to the size of the animal in younger individuals, because studies on checkerboard backgrounds have shown that in order to elicit disruptive patterns on checkerboards the size of a checker must exceed 40% of the size of the animal white square and have a contrast larger than 0.54, whereas our backgrounds had sand grain sizes below 20% of white square size and a coefficient of variance less than 10%, values fully consistent with a clearly uniform body pattern on checkerboard backgrounds (5).

A possible solution to the apparent paradox lies in considering alternative modes of camouflage. Age-dependent expression of disruptive components represents a clear bias from the point of view of matching background visual statistics, but it represents an adaptive behavior if we interpret the behavior of young cuttlefish as a form of masquerade (25-27). Disruptive components of the cuttlefish have often drawn comparison to pebbles found on the seabed. Thus, by expressing disruptive components on part of their body while adapting the intensity of the rest of their body to the background, young cuttlefish bear a close resemblance to a pebble on the seabed. Why would masquerade behavior decrease with age? Masquerade has been found to be more effective at deceiving predators when the inanimate objects that the camouflaging animals resemble are abundant in the environment (26). The statistics of pebble sizes in natural environments demonstrate a decreasing abundance of exemplars with increasing pebble size (28). Masquerade is thus predicted to be more effective in younger animals simply because smaller pebbles outnumber larger pebbles in most environments making masquerade more likely to be effective in deceiving predators in smaller animals.

## Conclusion

Our studies uncovered results that prompt a re-examination of several commonly held views about the rules of cuttlefish camouflage. Our finding of systematic intensity and color biases and the use of disruptive components on uniform backgrounds by younger animals all demonstrate that the hypothesis of optimal matching of background statistics alone is insufficient in explaining observed responses. Perceptual limitations, motor system dynamic range, alternative modes of camouflage and expected statistics (based on evolutionary knowledge) rather than actually observed statistics of the environment may need to be considered to explain animal responses on naturalistic backgrounds. Also, our work highlights problems in drawing conclusions across different modes of chromatophore-based behavior. Although intensity transients produced for the purpose of camouflage can be rapid (less than 1 second in duration) as has been found in the context of social communication and dynamic wave displays, the typical adaptation to background proceeds gradually over a timescale of several minutes.

## Methods

### Intensity matching protocol

Cuttlefish eggs collected from the North Sea were reared in tanks filled with recirculating artificial seawater until tested at 1 or 3 months of age respectively. Each animal was tested on all backgrounds. Cuttlefish were placed into a rectangular tank whose bottom was uniformly covered with a layer of loose sand. The thickness of the sand layer was around half a centimeter, sufficiently thick to encourage the animals to settle at the bottom of the tank and dig into the sand, but not so thick as to allow the animals to dig in to the point where their bodies would be covered by sand and obscured from view. The illumination of the bottom of the tank was uniform (less than 10% coefficient of variation) and constant for all conditions. The lamps were placed outside the tanks below the height of the water surface to avoid obscuring the image with reflections. The animals were recorded with a Sony camera acquiring images at 3 frames per second. After placement in the experimental tank, the animals were left undisturbed in the experimental room for 30 minutes. To determine the lightest intensity five animals were anesthetized in a magnesium chloride solution isotonic to artificial seawater and then each animal was observed at six random positions in the experimental tank containing white sand based on which the intensity was estimated.

### Image analysis

All image analysis was performed using custom-written code in Matlab. The sensitivity of the camera to object luminance was determined from a calibration curve to approximately follow a power law with an exponent of 0.65. Such a power law demonstrates diminishing sensitivity to changes with increasing intensity reminiscent of the luminance sensitivity of animal visual systems (which generally obey logarithmic sensitivity). We thus elected to present all our intensity measurements in camera units unless otherwise noted. For analysis of intensity matching, animals were manually segmented from the background and their mean intensity was then calculated. For analysis of adaptation time course, the difference between the red and blue image channels was calculated and low pass filtered with a 50*50 square kernel and the resulting image histogram subjected to kmeans clustering to detect the animal. The center of mass and mean intensity were calculated for each frame. Velocity was inferred from the motion of the center of mass. The algorithm was tested and produced reliable segmentation on the four darkest backgrounds.

For analysis of disruptive components a library of 170 images of sepia patterns was assembled. Each cuttlefish was segmented from the background, initially morphed so that the length and width of the mantle were equal for all images. 16 reliably identifiable reference points (see supplementary Figure 1) were then marked out on each image, from which a Voronoi tessellation was assembled and each triangle was then subjected to an affine transformation followed by bilinear interpolation to generate perfect match between target and template. The morphed library was subjected to hierarchical clustering using the built-in Matlab routine hclust using a Pearson correlation based distance metric. One-dimensional spectra of juvenile cuttlefish were calculated by morphing all animals to have the same width and height, Fourier transforming the images, binning 2D Fourier transform coefficients according to the modulus of their x and y frequency and summing the moduli of the coefficients within a bin to yield the frequency-energy distribution.

**Figure S1:**
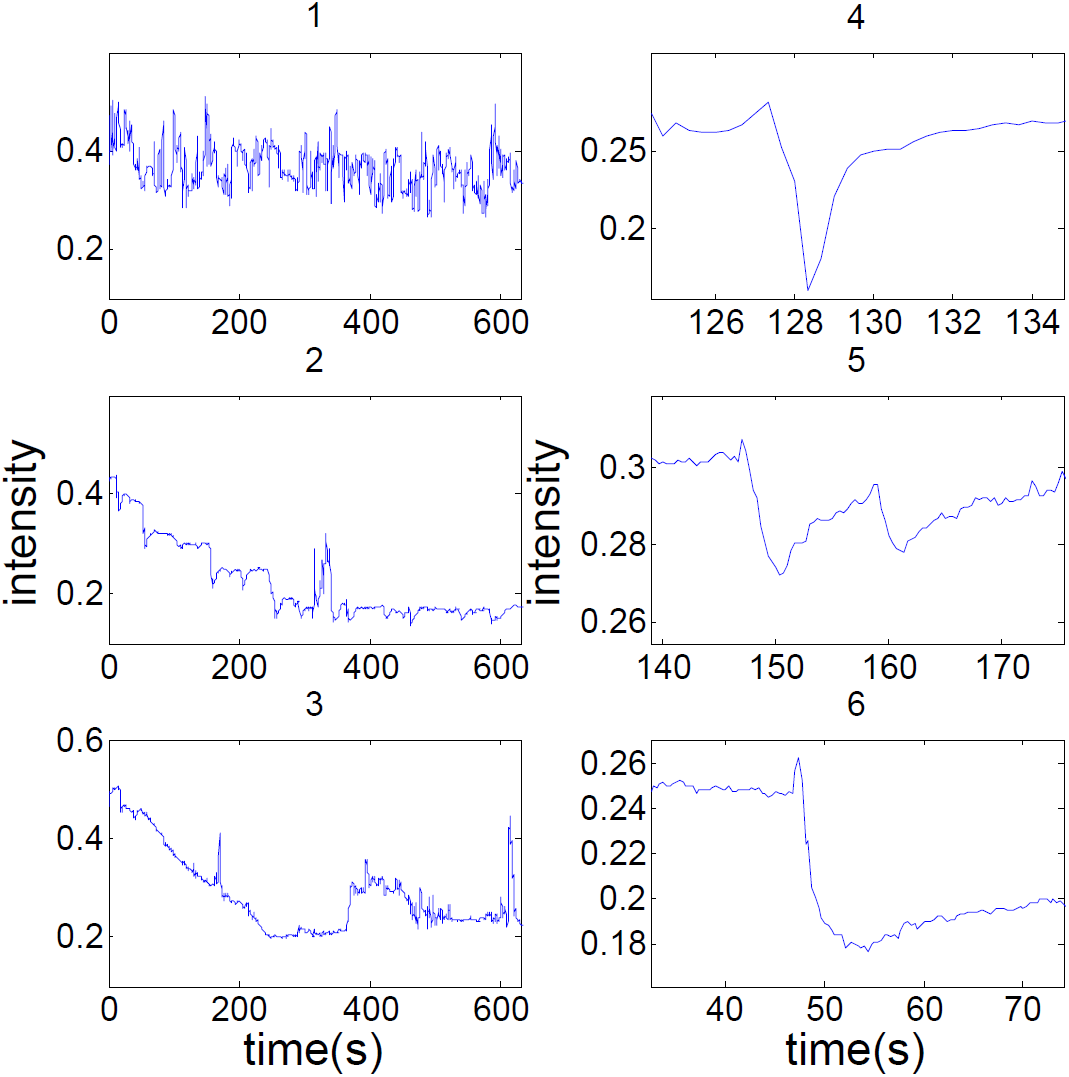
example traces showing diversity in individual animals’ responses when adapting to the background. 1: no adaptation; 2: adaptation in jumps; 3: slow linear decay. 4-6: zooms into local intensity transients, showing fast (4) to slow (6) processes.

## Acknowledgements

This work was funded by the Max Planck Society. A.L. performed these experiments as part of the IMPRS doctoral program. Michael Kuba provided advice on the zoology and handling of the animals throughout the project. Gilles Laurent managed the lab space in which the project was carried out. I thank the animal house stuff for assistance with keeping the animals.

## References

1. Messenger JB (2001) Cephalopod chromatophores: neurobiology and natural history. Biological Reviews of the Cambridge Philosophical Society 76(4): 473–528

2. Hanlon RT et.al. (2009)Cephalopod dynamic camouflage: bridging the continuum between background matching and disruptive coloration. Philos. Trans. R. Soc. Lond. B Biol. Sci. 364(1516): 429–437.

3. Caro T, Izzo A, Reiner RC, Walker H, Stankowich T (2013) The function of zebra stripes. Nat. Comm. 5 (3535)

4. Mills LS et.al. (2013) Camouflage mismatch in seasonal coat color due to decreased snow duration. PNAS 118(10): 7360–7365

5. Barbosa A et.al. (2008) Cuttlefish camouflage: The effects of substrate contrast and size in evoking uniform, mottle or disruptive body patterns. Vision res. 48(10): 1242–1253

6. Chiao CC et.al. (2013) How visual edge features influence cuttlefish camouflage patterning. Vision res. 83: 40–47

7. Chiao CC, Chubb C, Hanlon RT (2007) Interactive effects of size, contrast, intensity and configuration of background objects in evoking disruptive camouflage in cuttlefish. Vision res. 47:2223–22358

8. Shashar N, Rutledge PS, Cronin TW (1996) Polarization vision in cuttlefish- a conclealed communication channel? J. Exp. Biol. 199: 2077–2084

9. Zylinsky S, Darmaillacq SA, Shashar N (2012) Visual interpolation for contour completion by the European cuttlefish (Sepia officinalis) and its use in dynamic camouflage. Proc. R. Soc. B. 279(1737): 2386–2390

10. Mathger LM, Barbosa A, Simon M, Hanlon RT (2006) Color blindness and contrast perception in cuttlefish (Sepia officinalis) determined by a visual sensorimotor assay. Vision research 46(11): 1746–1753

11. Hanlon RT, Messenger JB (1988) Adaptive coloration in young cuttlefish (Sepia officinalis L.): the morphology and development of body patterns and their relation to behavior. Philos. Trans. R. Soc. Lond. B Biol. Sci. 320(1200): 437–487

12. Kelman EJ, Osorio D, Baddeley RJ (2008) A review of cuttlefish camouflage and object recognition and evidence for depth perception. J. Exp. Biol. 211: 1757–1763

13. Julesz B (1981) Textons, the elements of texture perception, and their interactions. Nature 290(5802):91–7

14. Portilla J, Simoncelli EP (2000) A parametric texture model based on joint statistics of complex wavelet coefficients. Int. J. Comput. Vision 40: 49–71

15. Ringach D (2004) Mapping receptive fields in primary visual cortex. J. Physiol. 558(3): 717–728

16. Seelig JD, Jayaraman V (2013) Feature detection and orientation tuning in the Drosophila central complex. Nature 503(7475):262–266

17. Messenger JB, Nixon M, Ryan KP (1985) Magnesium Chloride as an anesthetic for cephalopods. Comp. Biochem. Physiol. 82(1):203–205

18. Zylinsky S, Osorio D, Shohet AJ (2009) Cuttlefish camouflage: context-dependent body pattern use during motion. Proc. Roy. Soc. B. 276(1675): 3963–3969

19. Laan A, Gutnick T, Kuba M, Laurent G (2014) Behavioral Analysis of Cuttlefish Traveling Waves and Its Implications for Neural Control. Curr. Biol. 24(15): 1737–1742

20. Malik J, Belongie S, Leung T, Shi J (2001) Contour and texture analysis for image segmentation. Int. J. Comput. Vision 43(1):7–27

21. Stuart-Fox D, Mousalli A (2009) Camouflage, communication and thermoregulation: lessons from colour changing organisms. Philos. Trans. R. Soc. Lond. B Biol. Sci. 364: 463–470

22. Norris KS (1967) Color adaptation in desert reptiles and its thermal relationships. In Lizard ecology—a symposium (ed. Milstead WW): 162–229. Columbia, MO: University of Missouri Press

23. Nise N (2010) Control systems engineering (6^th^ ed.)

24. Kittle C, Kroemer H (1980) Thermal Physics (2^nd^ ed.)

25. Skelhorn J, Rowland HM, Ruxton GD (2010) The evolution and ecology of masquerade. Biological Journal of the Linnean Society 99: 1–8

26. Skelhorn J, Rowland HM, Speed MP, Ruxton GD (2010) Masquerade: camouflage without crypsis. Science 327(5961):51

27. Buresch KC et.al. (2011) The use of background matching vs. masquerade for camouflage in cuttlefish Sepia officinalis. Vision res. 51(23-24):2362–2368

28. usSeaBed: Coastal and marine geology program

